# Neural activity in a hippocampus-like region of the teleost pallium are associated with navigation and active sensing

**DOI:** 10.1101/495887

**Authors:** H Fotowat, C Lee, JJ Jun, L Maler

## Abstract

Neural mechanisms underlying spatial navigation in fish are unknown and little is known, for any vertebrate, about the relationship between active sensing and the formation of spatial maps. The weakly electric fish, Gymnotus Carapo, uses their active electric sense for spatial navigation. The electric organ discharge rate (EODr) undergoes transient increases during navigation to enhance electrosensory sampling. Gymnotus also uses stereotyped forward/ backward swimming as a second form of active sensing that brings objects towards the electroreceptor-dense head region. We wirelessly recorded neural activity from the pallium of freely swimming Gymnotus. Spiking activity was sparse and occurred only during swimming. Notably, some units exhibited significant place specificity and/or association with both forms of active sensing. Our results provide the first characterization of neural activity in a hippocampal-like region of a teleost fish brain and connects active sensing via sensory sampling rate and directed movements to higher order encoding of spatial information.

## Introduction

Neural mechanisms underlying spatial navigation has been intensely studied for decades in mammals and, especially, in rodents (Eichenbaum 2017, Moser et al 2017, Chersi and Burgess 2015). Various cell types in different areas of the hippocampal formation have been identified to encode various self, environmental, and social cues that enable the animal to successfully navigate towards a goal. These cells include place cells (O’Keefe and Dostrovsky 1971), boundary cells (Saveli et al 2008, Solstad et al 2008), grid cells (Saveli et al 2017), head direction cells (Taub 2007), goal direction cells (Sarel et al 2017), and more recently social place cells (Danjo et al 2018, Omer et al 2018).

Remarkably, there is a large degree of similarity in the neural representation of space among phylogenetically distant mammals (e.g. between echolocating bats and rodents: Ulanovsky and Moss 2007). It is known that teleost fish, whose common ancestor with mammals lived approximately 450 million years ago are also capable of spatial learning (Rodriguez et al 2002, Jun et al 2016). Whether these fish utilize the same neural mechanisms for spatial learning as mammals do remains completely unknown. In goldfish, studies based on lesion and cytochrome oxidase histochemistry have identified the dorsolateral pallium (DL) as selectively essential for spatial learning (Ocana et al 2017, Uceda et al 2015, Rodriguez 2002, Duran et al 2010, Broglio et al 2010). Based on these studies as well as patterns of connectivity and gene expression, it has been hypothesized that DL has similar connectivity and perhaps is even homologous to the mammalian hippocampus (Elliot et al 2017; Harvey-Girard et al 2012). Although one study reported recordings from one putative place cell near the medial edge of DL of gold fish (Canfield and Mizumori 2004), to our knowledge no studies have quantitatively demonstrated the presence of place-associated cells in the teleost pallium.

We chose to study the neural activity of the pallium of pulse-type weakly electric fish *Gymnotus sp*. in the context of spatial navigation. Through detailed analysis of their spatial learning behavior, Jun et al (2016) have shown that these fish can learn the location of food relative to landmarks in complete darkness relying mainly on their short-range active electrosensory system. This study further demonstrated that, in the process of learning, *Gymnotus* use several active sensing strategies. These include increasing the rate of their electric organ discharge (EOD), which results in an increased rate of sensory sampling, as well as back and forth swimming (B-scans) past landmarks (Jun et al 2016). These fish therefore provide an excellent model system for reading out the dynamics of sensory sampling and “attentive state” and relating this information to neuronal activity associated with landmarks during spatial navigation. Interestingly, once the fish have learned about the location of food, they appear to identify landmarks they encounter along their current trajectory, and then move towards the food location despite not being able to electrosense other landmarks or food from afar. Based on this finding, Jun et al (2016) hypothesized that fish’s trajectories from a learned landmark to food location were based on path integration.

Changes in electric field caused by the presence of landmarks are sensed by thousands of electroreceptors located on the fish’s skin. Primary afferents relay this information to the electrosensory lateral line lobe which then projects to the mid-brain torus semicircularis that provides electrosensory input to tectum (Krahe and Maler 2014, Carr et al 1982). Cells in the fish’s tectum respond to both electrosensory and visual object motion (Bastian 1982) and would therefore be expected to discharge as the fish swims past landmarks. The tectal cells responsive to object motion project to the preglomerular nucleus (PG), an analog or homolog of the mammalian thalamus (Giassi et al 2012b; Wallach et al. 2018), which in turn projects to DL. Based on its connectivity and gene expression, DL has characteristics of both mammalian cortex and the hippocampus (Elliott et al 2017). DL projects massively and in a highly convergent manner to the intermediate subdivision of dorsal pallium (DDi); DDi then provides strong feedback to DL via a magnocellular component of DD (DDmg, Giassi et al 2012c, Trinh et al 2016, Elliott et al 2017). DL has vastly more neurons than either DDi or DDmg (Supplementary Figure 1, Trinh et al 2016), and they are far more densely packed (Giassi et al 2012c). We therefore decided that sampling DDi would be far more efficient than sampling DL and would still give us an idea of the kind of spatial information that might be extracted by the teleost pallium.

A recent paper has suggested that DDi has similar connectivity, and might be homologous to, the mammalian CA3 region and might therefore be expected to contain cells responsive to spatial and/or motion features of the environment (Elliott et al 2017). We therefore used a wireless transmitter system to record neural activity from DDi as the fish swam freely in the dark in an experimental tank (open maze) containing differently shaped landmarks – we used the same tank and landmarks as in the Jun et al (2016) study so that we could directly compare behavioral and physiological data. We simultaneously tracked the fish’s position and recorded its EOD signal using electrodes placed inside the tank (Jun et al 2014a). We asked: what are the neural dynamics of cells in DDi and how do they relate to the rate of sensory sampling, the fish’s location and scanning movements relative to landmarks.

## Results

### Spikes were sparse and occurred mainly during movement

Figure 1 shows the wireless transmitter, the tetrode and their assembly prior to (A-C), and after implantation (D). Extracellular recordings from neurons within the dorsal pallium (DD) were wirelessly transmitted as the fish freely swam in a large circular tank that contained differently shaped landmarks (Figure 1E, see Methods). We aimed our tetrodes to the intermediate subdivision of DD: DDi (see Methods, Supplementary Figure 1). Figure 2A shows an example extracellular recording together with the raster plot for the four units that could be sorted based on their shape. Figure 2B shows the shape of the sorted units and their first three principle components that were used for clustering units (see Methods). The EOD rate (bottom red trace, Fig. 2A) was calculated using the EOD signal that was simultaneously recorded by the tank electrodes (see Methods). Figure 3A shows the distribution of the average firing rates of all units recorded from all the fish and across all trials, highlighting sparsity of spiking in this region (25 units, 5 fish, 23 trials, 20 hours and 36 min of recording). We found that that units in DDi fired at strikingly low rates, and that it was very unlikely to encounter a spike at low swimming speeds or during periods of quiescence. Figure 3B shows the histogram of swim speeds calculated across all time bins and trials (purple) and those in time bins where there was a spike (green). Notice a peak at low swim speeds in the purple histogram, which is missing in the green one. It is known that at very low speeds, that is during moments of behavioral quiescence or “down-states”, the EOD rate is lower than when the fish is active (generally less than 50 Hz down states in *Gymnotus sp*., Jun et al 2014b), resulting in a reduction in sensory sampling rate. Consistently, we found that spikes were highly unlikely during down-states (Figure 3C, notice the absence of spikes at EODr < 50 Hz) indicating that spikes in this region are likely linked to both sensory and motor activity.

**Fig 1.**
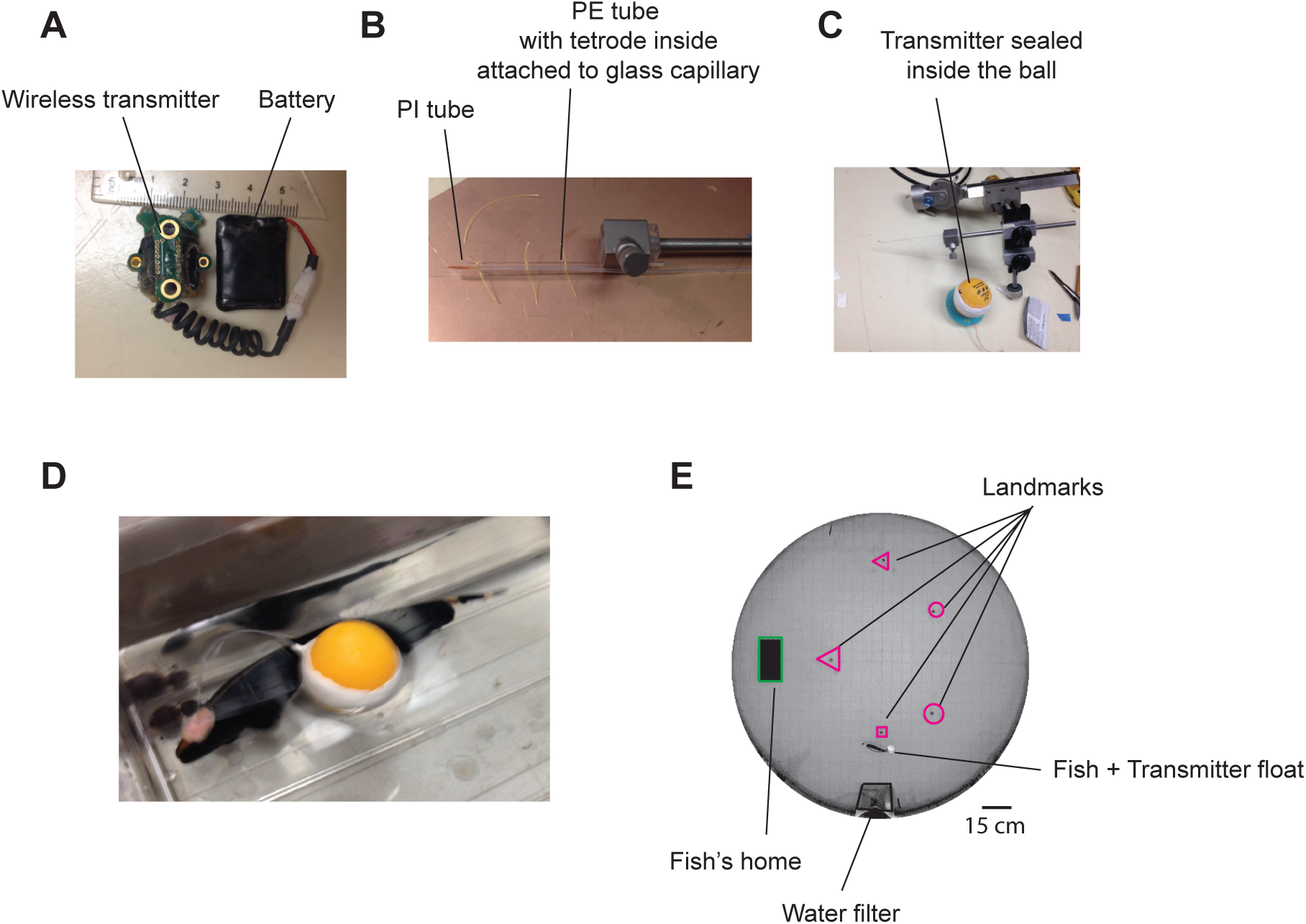
Experimental setup and example wireless recording. A. Wireless transmitter system used for recordings. B. A long tetrode was constructed and mounted on an electrode holder to be attached to a micromanipulator. C. The other end of the tetrode was connected to the transmitter, and the ensemble was placed in a ping-pong ball and sealed. D. Pictures of a fish after tetrode implantation together with the transmitter float (picture from a related species with the same size). E. Recordings were performed in a large experimental tank containing various landmarks made with clear and opaque plexiglass, as well as a water filtering system.

**Fig 2.**
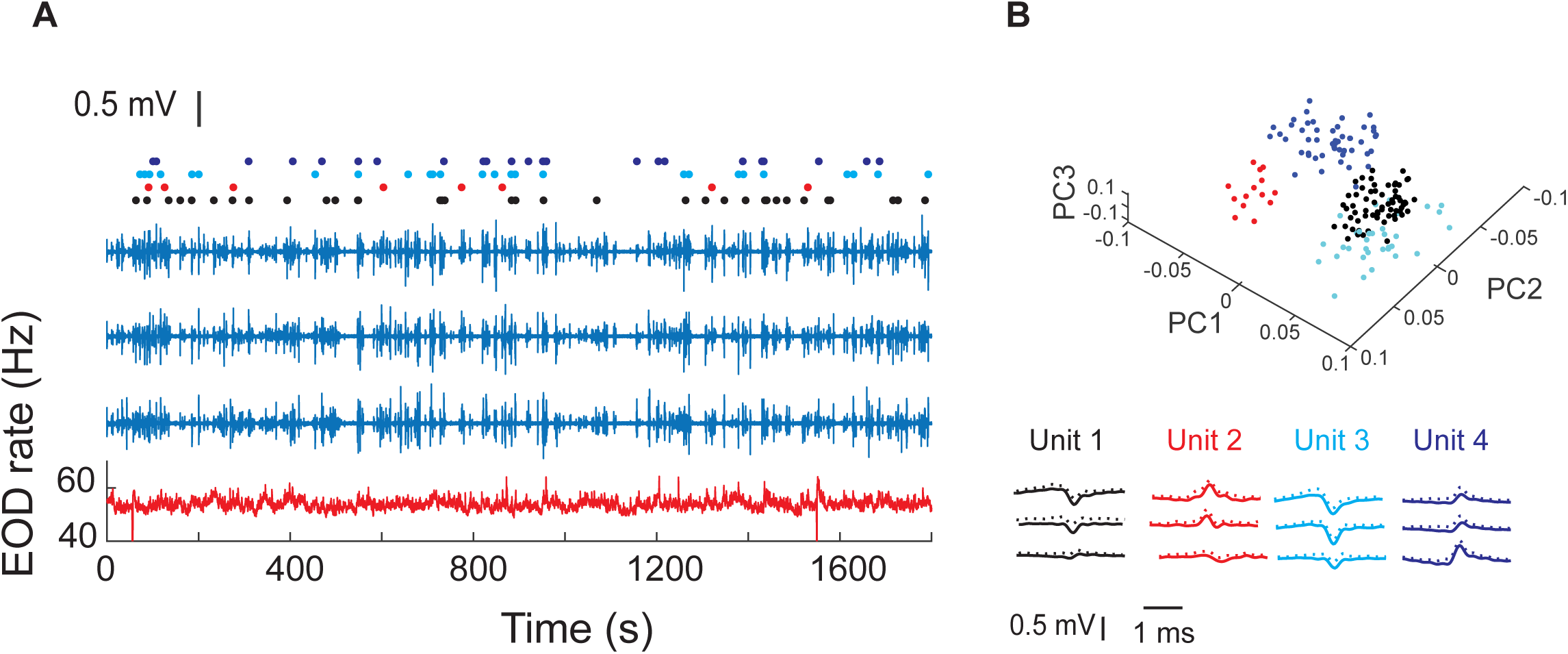
Example recordings in one fish and four isolated units. A. The blue traces show extracellular recordings on the three of the four tetrode channels after EOD spike removal (see Methods). The fourth channel was used as reference. The red trace shows instantaneous EOD rate calculated based on EOD recordings obtained by electrodes inside the experimental tank. Raster plots show spike timing corresponding to the four units. B. Average (SD) of waveforms of the isolated units from this recording and their first three principle components (PCs).

**Fig 3.**
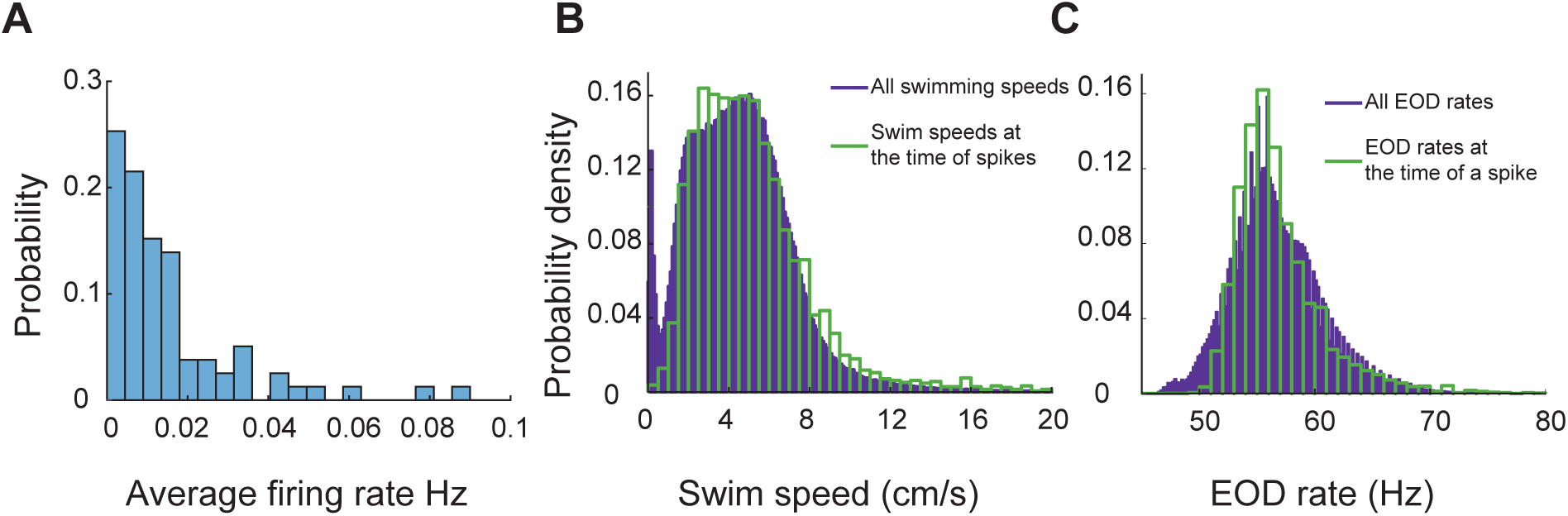
DDi units spiked sparsely during periods swimming and active sensing. A. Probability histogram of average firing rates observed in all units and all trials (25 units, 23 trials, 5 fish). B. Probability density function of all observed swim speeds (purple bars), and swim speeds at the time of a spikes (green bars). Spikes were unlikely at very low swimming speeds. C. Probability density function of all observed EOD rates (purple bars) and those observed at the time of spikes (green bars). Spikes were unlikely at low EOD rates (<50 Hz) corresponding to down states.

### Relation between spiking activity, EOD rate, swim speed and sampling density

The electric organ discharge rate (EODr) continuously varies in time (Figure 2, red trace). Higher EODrs indicate higher sensory acquisition rates, and large, fast transients in the EODr has been previously reported to occur near landmarks in the context of spatial learning (Jun et al 2016). Fish swim speed is variable and is also associated with proximity to landmarks and changes during learning (Jun et al 2016). Sampling density, the number of EOD pulses per unit length, depends on both EODr and swim speed and therefore also varies strongly with the fish’s location near landmarks (Jun et al 2016). Below, we describe the relation between DDi spiking and EODr, swim speed and, consequently, with sampling density.

*EODr:* To examine the relation between sensory acquisition rate and spiking activity of DDi units we calculated spike-triggered-EODr (stEODr) averages for all units that fired more than 10 spikes in a trial (see example in Figure 4A). We found that for the majority of units (15 out of 21), the mean EODr in an 8 second window around the spike time was significantly higher than that calculated for randomly time-shifted spikes (Fig 4A black trace). Moreover, we found that for many units the mean EODr increased prior to spike time and peaked within a few hundred milliseconds around the time of the spike. 13 out of the 21 units showed a significant peak in the stEODr average.

**Fig 4.**
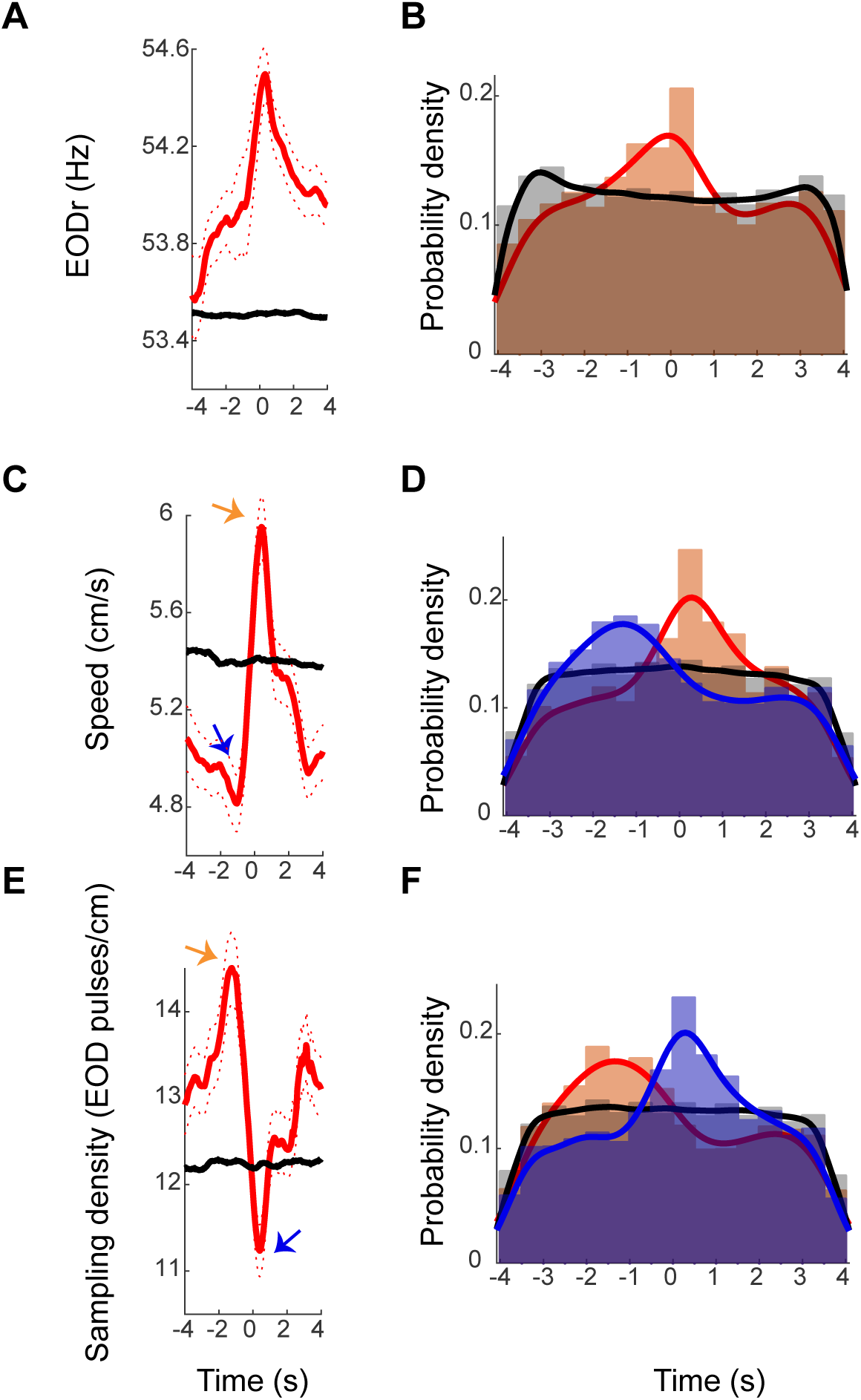
Examples stEODr, stSpeed and stSmpD averages. A. StEODr average for an exemplar unit (solid red curve, dotted curve: standard error). The black curve shows the stEODr average for 100 random time shifts of the same spike train. The dashed vertical line corresponds to the spike time (zero). B. Probability density of timing of stEODr peaks around individual spikes for units with significant peak in stEODr average (orange bars, 13 units in 5 fish) and for 100 random time shifts of the same spike trains (grey bars). There was a clear peak of stEODr around zero that did not exist in the random data. C. StSpeed average (red curve) for the same unit shown in A, and the corresponding average for randomly shifted spike times (black curve). Arrows point to the dip and peak in the stSpeed pre- and post-spike respectively. D. Probability density of speed dips (blue) preceding the spike and peaks (orange) after spikes for units with a significant peak in stSpeed average (14 units in 5 fish, grey: randomly time-shifted spike trains). E. stSmpD average for the same unit. stSmpD showed a clear peak before the spike and sharply decreased immediately post-spike, with the minimum occurring after spike time. F. Probability distribution of timing of the peak (orange) and dip (blue) in sampling density for individual spikes of units with significant peaks in both stEODr and stSpeed shows the same pattern of a pre-spike increase followed by a post-spike decrease (10 units in 5 fish). Fits in all histograms are non-parametric fits with Gaussian kernels.

Figure 4A shows an example unit with a significant peak in its stEODr average (stEODr for all the other units are presented in Supp. Fig 2). Figure 4B shows the probability density histogram of timing of peaks in EODr relative to individual spikes from all units and all fish (orange bars). The peaks were more likely to occur around the timing of the spike; within a second prior to and 0.5 s after spike time (median peak time was zero). The histogram for the timing of stEODr calculated using randomly time-shifted spike trains did not show such a peak around the spike time (grey bars).

*Swim speed*: We next considered the correlation between swim speed and spiking of the units. Interestingly, we observed that the fish’s spike-triggered-swim-speed (stSpeed) often decreased to a minimum (dip) before the time of spike, followed by a peak after (Figure 4C, blue and orange arrows, respectively. see also Supp Fig 2). 14 out of 21 units showed a significant peak in stSpeed average after the spike time. 10 out of these 14 units also had a significant peak in their stEODr average, with all stEODr average peaks except one occurring after the spike time (Supp. Fig 2, Fish 5, 1^st^ unit). Figure 4D shows the distribution of the timing of speed dips (blue bars) and peaks (orange bars) across all units and fish (median stSpeed dip time =-0.56 s median stSpeed peak time = 0.2 s). Unlike the stEODr, the timing of the dips and peaks in stSpeed were significantly shifted to negative and positive values i.e. before and after the timing of the spike, respectively (for dips: p= 4.4 e-24, for peaks: p=8.5e-11, non-parametric sign test). This pattern was not evident for the timing of peaks and dips of stSpeed calculated for randomly time-shifted spikes (grey bars show the peak times, similar pattern was observed for the dips, data not shown).

*Sampling density:* We next used units that showed significant peaks in both stEODr and stSpeed averages (10 units) and calculated the spike-triggered sampling density as the ratio of stEODr to stSpeed (stSmpD, exemplar unit is shown in Figure 4E, see Supp. Fig 2 for all units). The sampling density has units of EOD pulses emitted per unit length (pulses/cm), it is a measure of electrosensory spatial acuity, and has been previously proposed to be an indicator of spatial attention (Jun et al 2016). Mirroring stSpeed, we found that for most units the stSmpD average showed a peak before spike time and a dip after it (Figure 4F orange and blue arrows, respectively). Figure 4F shows the probability distribution for the peaks (orange bars) and dips in stSmpD (blue bars) for all units and all fish (median stSmpD peak time =-0.56 s, median stSmpD dip time = 0.24 s). The timing of the peak and dip in stSmpD were not significantly different from the timing of dip and peak in stSpeed, and occurred significantly before and after spike time, respectively (for peaks: p= 2.68 e-24, for dips: 1.1 e-10, non-parametric sign test). This pattern was not present in the timing of dips and peaks calculated for stSmpD of randomly time-shifted spikes (grey bars show the peak times, similar pattern for the dips, data not shown).

In summary, despite a large degree of variability in the dynamics of EODr and speed around individual spikes as indicated by the width of the timing histograms, we found that DDi spikes were most likely to occur after a dip in swim speed. EODr often increased prior to a spike and peaked around the time of the spike. The combination of the pre-spike decreasing swim speed and increasing EODr resulted in a strong increase in SmpD before the spike. Similarly, the post-spike increasing swim speed and decreasing EODr resulted in a strong SmpD dip post-spike. The changes in swim speed were clearly the dominant factor contributing to this change in SmpD. This is completely consistent with the results of Jun et al (2016), which shows that sampling density increases in the vicinity of landmarks. The net effect of these coordinated changes in EODr and swim speed leads to strongly enhanced electrosensory sampling (SmpD) over the trajectory traversed ~1-3 s before a spike followed by reduced sampling for up to 3 s after the occurrence of the spike.

### Spiking occurred near boundaries and landmarks

Units in DDi tended to fire in multiple locations within the experimental tank often close to the tank boundaries and landmarks. Figure 5 shows examples of swimming trajectories, spiking locations, and firing rate maps for 8 units recorded in 5 fish. To quantify place-specificity of each unit’s activity we calculated the “place information” in bits/spike (see Methods) for the 21 units that fired more than 10 spikes during the trial. The average of information across all units and trials was 1.58 bits/spike (SD= 1.15). For each unit and trial, we tested the statistical significance of place information level by comparing it to that calculated from randomly time-shifting the spike train relative to swim trajectory. Examples of the firing pattern of units that conveyed significant place information, and those that did not are given in Figure 5A and B, respectively. 11 of the 21 recorded units in 4 fish (3,3,3,2 units in each fish respectively) showed significant place specificity. Interestingly, 8 out of 11 units with significant place selectivity also had significant peaks in their stEODr averages (Supplementary Figures 2 and 3). Four of these units (2 fish) further showed a clear peak in SmpD before spike time. There was no clear correlation between lack of place specificity and absence of a peak in stEODr (5 out of the 10 units that were not place specific, also showed a significant peak in their stEODr average, and the other 5 did not, Supplementary Figure 3).

**Fig 5.**
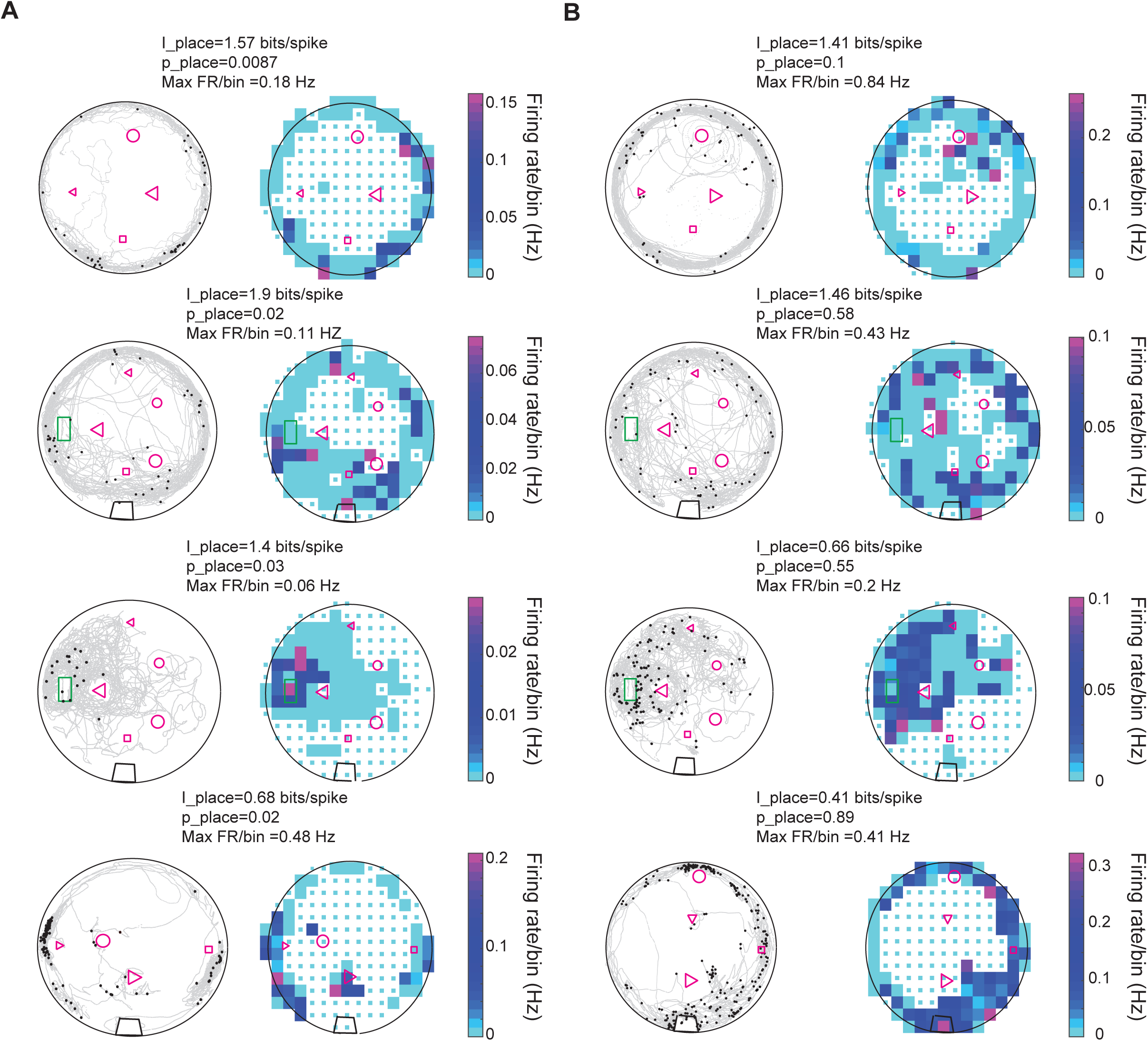
Spatial properties of DDi units. A. Left columns: Examples of spiking patterns of four units in four fish that conveyed significant place information. Grey curves show fish’s swimming trajectory, and black dots represent spikes. Right column: Firing rate maps of the same units shown to the left. Firing rate per bin is calculated by dividing the total number of spikes by the time spent in that bin. Only bins where the fish visited more than 5 times are used for the plot. The range of the color plot was clipped to the 97th percentile of the data for visual representation purposes (see Methods). The value of the maximum firing rate/ bin (Max FR/bin) is indicated above the plot. The place information for the unit and its level of significance compared to randomly time shifted spike trains are indicated as I_place and p_place, respectively. B. Same as A but for units that did not show statistically significant place specificity. The green rectangle corresponds to fish’s home, the black trapezoid in the bottom three plots show the location of a water filter. Other shapes represent various landmarks placed in the tank. Note that the fish could go inside the home area, but not other landmarks. Small cyan squares denote bins excluded from analysis due to visit counts less than 5 (see Methods).

Interestingly, changes in landmark configurations often resulted in changes in the firing patterns of the units and gain or loss of significance in conveyed place information. Figure 6A shows examples of the effect of changes in landmark configuration on spiking activity of three units. Figure 6B shows a summary plot of the effect of landmark removal on the firing rate of the units in a 10 cm radius around the location of the landmark (16 trials and 10 units, 12 out of 16 trials showed a reduction). For example, removing landmarks often resulted in a reduction in firing rate of the unit around the location of the missing landmark. Remarkably, in one case where the landmark was moved to a new location, the unit shifted its preferred firing rate to the new location of that same landmark (Figure 6A-iii, small triangle). Further, comparison of the firing rates in the two conditions normalized to the largest rate showed that the rate was significantly reduced with landmark removal (Figure 6B, inset). Therefore, we conclude that a subset of DDi units fire in a location-specific manner, and that their spatial specificity is tightly linked to the presence of landmarks.

**Fig 6.**
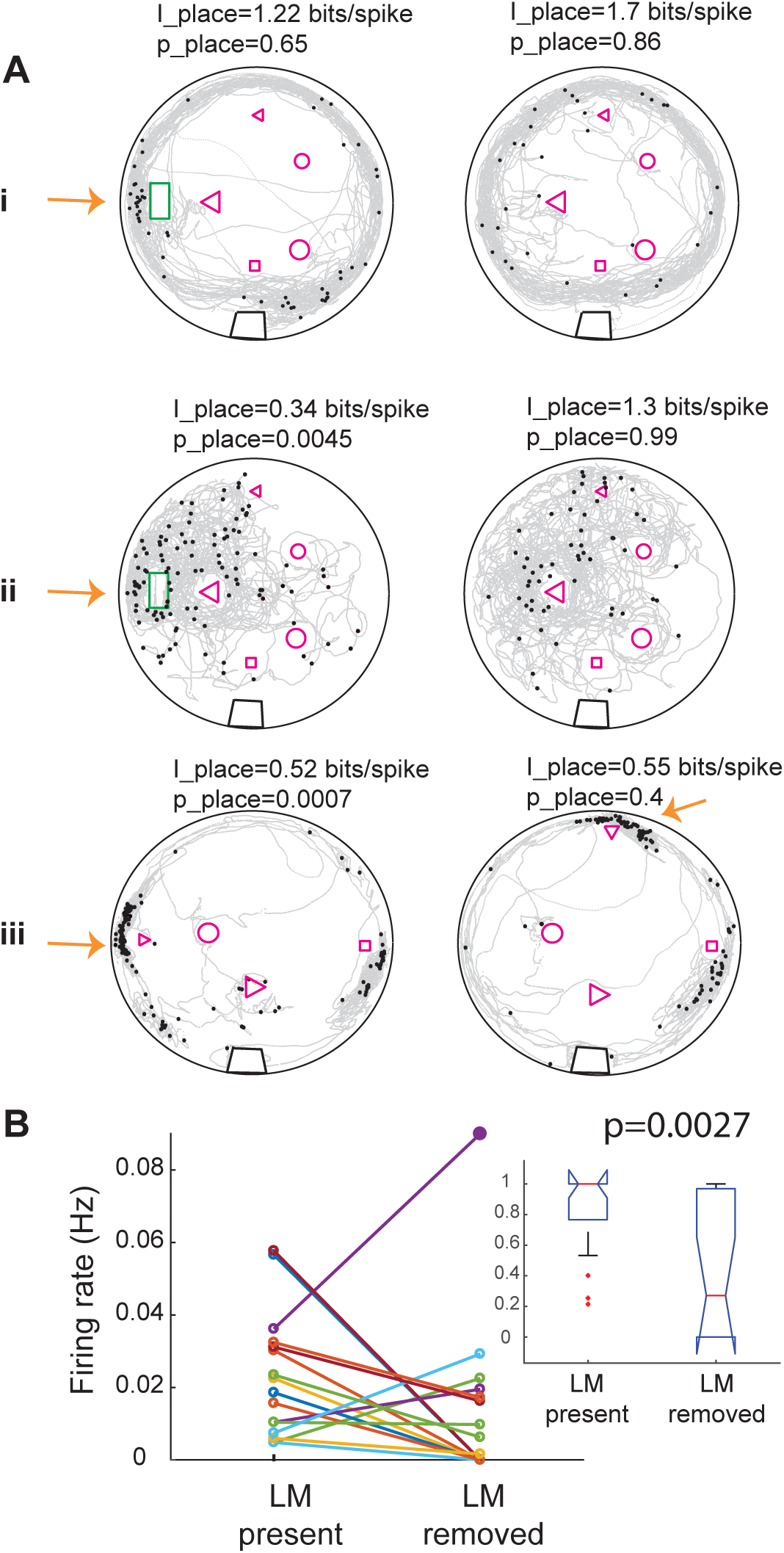
Removing landmarks often resulted in reduction of firing rate near the missing landmark. A. Examples of the effect of removing or displacing a landmark in three units in three fish. In i and ii the home was removed and in iii the small triangle was moved to a new location. Note that spikes now occur near the triangle in its new location. Place information and its significance level compared to randomly time-shifted spikes are shown above each panel. Arrows point to the location of the moved landmark. B. Summary plot of the effect of landmark removal on the firing rate of 10 units in 3 fish, 16 trials. In 12 out of 16 trials the firing rate decreased near the removed landmark. Inset: comparison of the normalized firing rates between the two conditions for the same data set. Firing rate was significantly lower in after landmarks were removed. Kruskal-Wallis p value is indicated on the plot.

### Swim direction and relative landmark location preference

*Gymnotus* can swim in both forward and backward directions. Backward swims are often observed in the context of prey capture and spatial navigation and serve to bring prey or landmarks that the fish has swam past, back near the fish’s head and rostral trunk (Pedraja et al 2018, Jun et al 2016, Nelson and Maciver 1999), the regions containing the highest density of electroreceptors (Carr et al 1982 and Castello et al 2000).

Interestingly, we found that most units in DDi showed preference for spiking during backward swims. Figure 7A shows the distribution of the swim direction preference indices (see Methods) for all units in all fish. The distribution showed a significant bias towards negative values that correspond to backward swims (mean (SD) =-0.196 (0.33), p=0.0072, non-parametric sign test).

**Fig 7.**
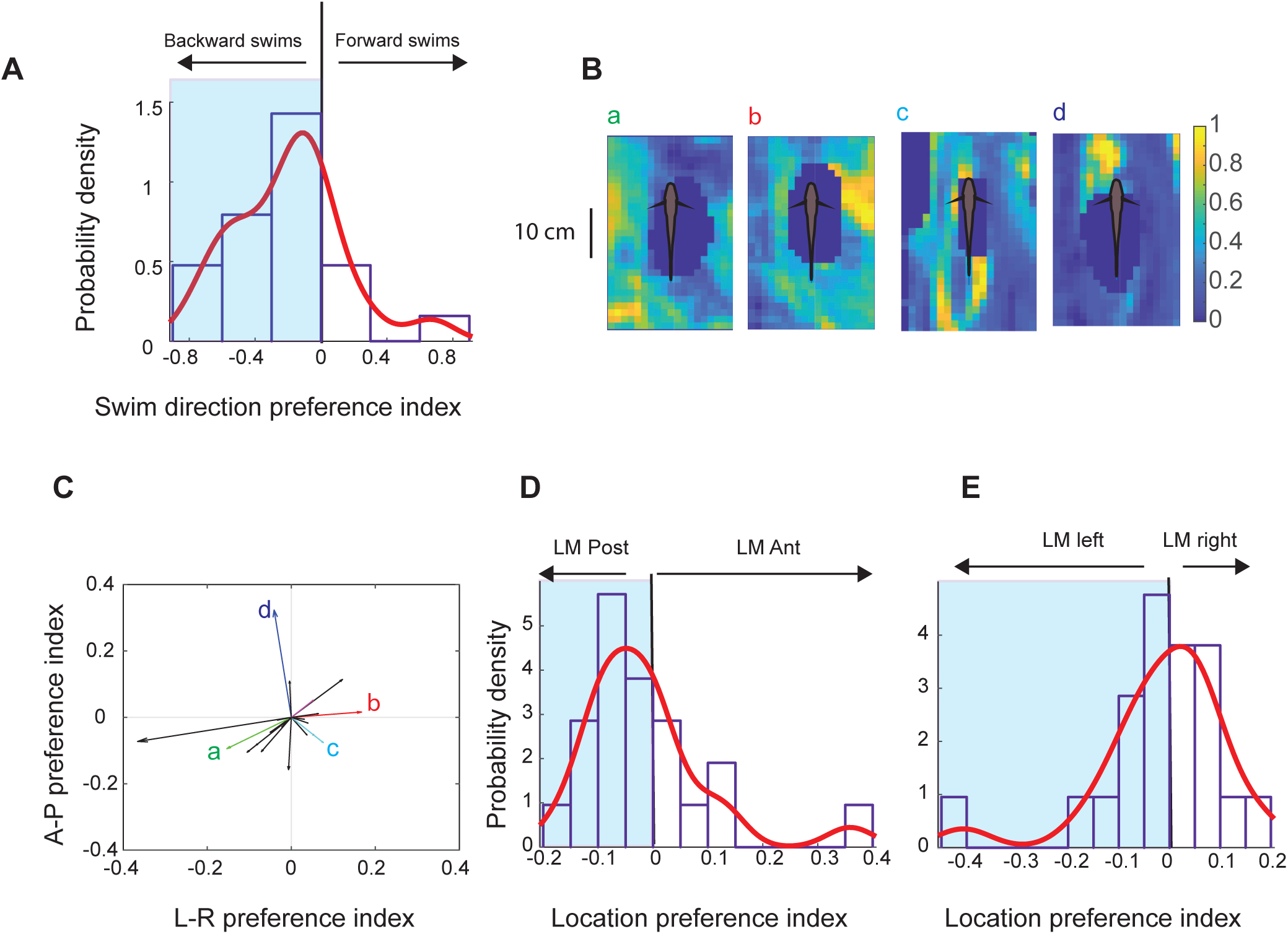
Relationship between swimming direction and preferred landmark location. A. Probability distribution of direction preference index (21 units, 5 fish). Negative values correspond to preference for spiking during backward swims, with the maxima +/-1 corresponding to spiking only during forward or backward swims, respectively. The red curve shows a non-parametric fit with Gaussian kernels. There was a significant bias for spiking for negative swim direction preference indices (p = 0.0072, non-parametric sign test). B. Examples of normalized landmark location probability. Units b and c are from the same fish and other units are from three other fish. C. Vector plot of the direction preference indices for anterior-posterior (AP) and left-right (LR) preference indices. Small letters next to the arrows correspond to exemplar units shown in panel B. D. Most units showed preference index for landmarks that were located on the posterior 2/3 of the body (blue shaded area). E. Left-right preference indices were equally likely to be positive or negative.

We next asked, do units in DDi have a preferred landmark location, i.e. are they more likely to fire when a landmark has a specific location relative to the body of the animal? To answer this question, we calculated a probability map for landmark locations when a given unit was active. To do this we divided spike-triggered landmark locations by position-triggered landmark locations (see Methods). Figure 7B shows normalized landmark probability at the time of spiking of four exemplar units. For each unit, we then calculated an anterior-posterior and a left/right preference index (see Methods). Figure 7C shows preference vectors plotted using these preference indices for all units in all fish. Vectors corresponding to exemplar units in Figure 7B are indicated within this plot. We found that for most units the probability of landmark presence was higher in the region posterior to the body of the fish (Figure 7D, 14 out of 21 units). This bias was not present for left-versus right location for the landmarks (Figure 7E, 10 out of 21 units preferred spiking to contralateral objects). Because in all fish the tetrode was implanted in the left lobe of the pallium, these results indicate that activity in DDi is not unilateral. In summary, most DDi units we recorded spiked in response to objects located in the region posterior to the body of the fish as fish was swimming backwards. These units therefore appear to correlate to a scanning movement that would lead to relocating these objects from the trunk to the head and more anterior parts of the body, which contain the highest density of electroreceptors.

## Discussion

In this study, we wirelessly recorded and characterized neural activity within the dorsal pallium (DDi) of freely swimming pulse-type weakly electric fish *Gymnotus sp*. This is the first study to report on location and movement related response properties of cells in a region of a teleost fish’s pallium that has similar connectivity, and may be homologous to, the CA3 area of mammalian hippocampus (Elliot et al 2017). A previous study (Elliott et al., 2015), using a related (immobilized) gymnotiform fish (*Apteronotus leptorhynchus*), found that, in the absence of sensory input, DDi cells were completely silent. In response to electrocommuncation and acoustic signals, DDi cells discharged very sparsely, at long latencies and with a small number of spikes; these results are comparable to our data. Further work will be required to determine whether different DDi cells respond to electrolocation versus electrocommunication signals.

Gymnotiform fish utilize two active sensing behaviors. Pulse type fish (e.g *Gymnotus* sp.) will increase their EODr and sampling density near landmarks and food (Jun et al 2016); *Apteronotus* is a ‘wave type’ fish and maintains a constant EODr when foraging. Both Gymnotus (this paper; Jun et al., 2016; Pedraja et al 2018) and *Apteronotus* (Nelson and Maciver 1999) will use back-and-forth scanning movements (B-scans) in proximity to salient landmarks.

In the discussion below we first summarize our most important results and then interpret them with respect to neuroanatomy (effectively identical in *Apteronotus* and *Gymnotus*, Giassi et al. 2012a-c) and the results presented in Jun et al (2016). Critical to our interpretation is that our experiments were carried out in naïve fish, i.e., animals that had not learned the spatial layout of the tank (open maze) environment. We therefore compare our data to the ‘early learning’ results of Jun et al, where the fish also first encountered landmarks in the same open maze environment and, by using active sensing, learned to identify landmarks and their spatial relation to food. We also incorporate the recent results of Wallach et al. (2018), who examined the response of *Apteronotus* PG neurons to looming/receding and longitudinal object motion. We note that forward object motion is equivalent to the backward swimming component of B-scans. Similarly, backward object motion is equivalent to the forward swim direction.

We found that DDi units only fired when the fish was moving and the EODr was high (Figure 3, Up state: Jun et al. 2014b). Discharge typically occurred at multiple locations distributed within the experimental tank and often near tank boundaries and landmarks. Spikes from about half of the units we recorded conveyed significant spatial information (Figure 5). The spatial pattern of spiking and its spatial specificity changed when the landmark configuration was altered (Figure 6A). We found that in the majority of units, spikes were linked to the presence of landmarks and that removing landmarks resulted in a reduction in the firing rate in the location of the missing landmark(Figure 6B). These findings demonstrate that cells in DDi encode the presence of landmarks or boundaries, albeit in a stochastic manner, as spikes were not fired at every instance the animal visited a given location.

Individual spikes of many DDi cells are precisely linked to the types of active sensing (EODr and sampling density increases and B-scans) that occur when the fish is learning the spatial layout of the open maze environment (Jun et al 2016). We found that over the whole recording session, for many units on average, the spike-triggered EODr average started increasing prior to spike time and continued till slightly after the spike (13 out of 21 units, Figure 4 A, B). In many cells (14 out of 21 units), the spike-triggered speed showed a dip prior to a spike and dramatically increased post-spike (Fig. 4C, D). This combination resulted in an increased sampling density before and during the spike followed by its reduction post-spike. Wallach et al (2018) report that many PG cells fire as an object approaches the fish (*Apteronotus*). We hypothesize that, in *Gymnotus*, the increased EODr and sampling density will also strongly drive PG spiking and therefore DL spiking activity. The only sensory input to DDi is from DL which leads us to conclude that DDi spikes will likely feedback (via DDmg) to DL while DL is activated by PG input. We further discuss this point below.

In the context of active sensing these fish exhibit stereotyped forward/backward swimming motions (B-scans) that are thought to be important for learning about landmarks in the context of spatial navigation (Pedraja et al 2018, Jun et al 2016). Likewise, during prey capture, the fish tend to swim backwards once they have passed a prey item to align the prey with the head region where the highest density of electroreceptors is found (Nelson and Maciver 1999). Jun et al (2016) demonstrated that EODr and sampling density are highest during the backward phase of B-scans suggesting that PG will be very strongly driven by this phase of B-scans. Remarkably, Wallach et al (2018) described PG units that responded strongly and specifically throughout the forward movement of an object, i.e., the same relative motion that would occur during backward swimming. Most appropriately, we found that most units we recorded (17 out of 21) showed a preference for spiking during the backward swim phase of B-scans (Figure 7A). Moreover, we found that most units (12 out of 17) with preference for spiking during backward swims spiked most when objects were initially located near the trunk region (Figure 7C, D). In other words, these units spiked during a movement that would result in the object ending up near the head – the region with the highest density of electroreceptors. We hypothesize that this PG activity drives DDi cells (via DL) during back-swimming. The DDi spikes will feedback (via DDmg) to DL while it still being activated by its strong ongoing PG input. Under these conditions there will be three temporally overlapping sources of excitatory synaptic input to DL cells: (1) PG spiking driven by the backward scanning of the landmark, (2) DL spiking driven by its local recurrent connectivity (Trinh et al 2016), and (3) feedback input from DDi (via DDmg). As we have previously noted, DL is likely the site for storage of spatial memory. We therefore further hypothesize that the spatial learning described by Jun et al is driven by synaptic plasticity at one or more of these synaptic inputs to DL neurons.

Finally, we pose the question: are the spatially-specific units we find in this hippocampus-like region of the fish brain functionally similar to the location/landmark responsive cells found in the mammalian hippocampus? There are five clear aspects of DDi cell discharge that suggest similarity to these cells. 1. Hippocampal place and/or grid cells exhibit sparse firing (Diamantakis et al 2016; Hainmuller et al 2018; Rolls 2015) and only during movement (Chen et al 2013; Winter et al 2015; Song et al 2005; Harvey et al 2018). 2. Like place cells, the spatially specific DDi cells discharge is closely tied to the presence of a landmark (Muller and Kubie 1993) and, like place cells, they will discharge upon an early encounter with a landmark (Alme et al 2014, Wilson and McNaughton 1993). 3. In a ‘large environment’, place cells will exhibit multiple place fields (Fenton et al 2008, Park et al 2011) much like the DDi cell discharge near multiple landmarks. 4. Place cell discharge is very variable across different visits to their preferred place field (Fenton and Muller 1998) much like the variability we observed in DDi cell discharge across visits to the same landmark. 5. Lastly, and most interestingly, discharge of these DDi units near landmarks is associated with active sensing, i.e., increased EODr and sampling density and scanning behavior (B-scans). Rodents use head-scans as ‘a spatially directed investigative behavior’ (Poulter et al 2018); there is a clear functional analogy between head-scans and B-scans. An important recent study showed that head-scans drive formation or strengthening of place field discharge of hippocampal cells (Monaco et al 2014). We hypothesize that, during B-scans, DDi feedback to DL will also drive the formation of place/object-associated discharge of DL cells, and that this is a key element of spatial learning in gymnotiform fish.

In blind rats, place cells can have place fields far from any landmark (Save et al 1998). Unfortunately, the fish hardly ever swam far from the tank boundary and landmarks and so we cannot determine whether this is also true for DDi cells. Further experiments with fish that have been extensively trained in the experimental tank and willing to traverse long empty spaces will be required to investigate this critical issue.

We propose that place and/or landmark-associated cells are not unique to the mammalian hippocampus but, instead, have evolutionary precursors stretching back 450 million years to the last common ancestor of mammals and teleosts fish (Jun et al 2014b). A critical objective of future experiments will be to improve our ability to record neural activity across distinct pallial regions (e.g., DD, DC and DL, see Giassi et al 2012b) in freely swimming fish and over many days of exploration and spatial learning. This will allow us to gain a better understanding of the core neural mechanisms underlying spatial learning by revealing commonalities and differences between the mechanisms responsible for memory guided navigation in the teleost versus mammalian brain.

## Methods

### Wireless transmitter and tetrode fabrication

A wireless transmitter/receiver system (Figure 1A, TBSI-W16, Triangular Biosystems Intl, Durham, NC) was used for transmitting and receiving neural recordings in freely swimming fish. Tetrodes were constructed using 12-micron stablohm wires (stablohm 650, California Fine Wire Company, Grover Beach, CA) and wound using a Neuralynx Tetrode Spinner (Neuralynx, Bozeman, MT). Each tetrode was made 15-cm long and passed through slightly shorter flexible polyethylene tubing (PE, PE10, 0.61 mm OD × 0.28 ID, Warner Instruments Corp, Hamden, CT) such that either end of the tetrode was sticking outside the tube (Figure 1B). The one end of the tetrode to be implanted was further passed through a 0.5 cm long 180-micron diameter polyimide tubing (PI, Microlumen, 068-I, Lot#24331) and fixed in place using a mixture of Krazy glue and dental cement (Jet denture repair powder, Lang Dental Mfg. Co Inc., Wheeling, IL) for additional rigidity (Figure 1B). The PE tubing containing the tetrode was attached to a glass capillary, which could be solidly inserted inside an electrode holder attached to a micro-manipulator (Figure 1C). Melted Polyethylene glycol (PEG) was used to attach the tubing to the glass capillary. At room temperature PEG solidifies and acts as a water-soluble glue. Once the electrode was implanted, the tubing could be separated from the glass capillary by gently pouring water on the solidified PEG. The other end of the tetrode protruding from the PE tubing was also stabilized inside the PE tubing using another piece of PI tubing and glue. Individual tetrode wires were then separated at that end and attached to the input ports of the electrode interface board (EIB) which could then be plugged in to the transmitter. One of the 4 electrodes was used as reference. At this point the electrodes were first electroplated using Neuralynx gold plating solution and then with a solution of Ethylene Dioxythiopene monomer (EDOT) and Polystyrene sulfonate (PSS) using nanoZ plating protocols and software (MultiChannel Sytems MCS GmbH, Germany). Individual tetrode impedances varied between 100-200 kOhms. The EIB was then attached to the transmitter-battery ensemble and they were then put inside a cut-open Ping-Pong ball, which was used as a float for the transmitter system. A few pieces of vibration absorbing gel (Z8006, Kyosho, Lake Forest, CA) were added inside the ball and around the transmitter system to prevent movement and vibrations that could result in transmission noise. The Ping-Pong ball was then closed back (Fig 1C) and water proofed using mouldable glue (Sugru, London, UK).

### Animal preparation and electrode implantation

All animal procedures were performed in accordance with the regulations of the animal care committee of the University of Ottawa. *Gymnotus* sp. of either sex were used for all experiments. Before implanting, tetrodes were first cleaned by submersion in 70% ethanol for 15 minutes and then in sterile saline solution (0.9% NaCl) for another 15 minutes to rinse off the ethanol. They were then further sterilized using UV illumination. Fish were anesthetized in a small container with tricane methanesulfonate (MS-222; Aqua Life, Syndel Laboratories) in tank water solution. They were then transferred to a holder outside of water where their head and body could be stabilized in preparation for surgery and electrode implantation. The MS-222 solution in oxygenated deionized (DI) water was continuously administered in this setup through a tube that was inserted in the mouth, and the fish’s body was covered with wet sponges and Kimwipes to protect the skin. Before opening the skull, it was first completely dried, then some crazy glue was applied on the contralateral side of the planned implant to make the skull surface rough. This procedure helped with the final closure step as the dental cement mixture could better adhere to the remaining pieces of roughed bone. Additionally, a few indentations were made using a dentist drill on the surface of the contralateral skull. These indentations served as extra attachment points as they were filled with the dental cement mixture at the closing stage. The dorsal pallium was exposed by a small craniotomy and the tetrode was micro-manipulated to the DD region.

DD of gymnotiform fish is divided into superficial (DDs), intermediate (DDi) and magnocellular (DDmg) sub-regions (Giassi et al 2012a). Elliott et al (2017) suggested that DDi, on the basis of its connectivity pattern, was similar and perhaps homologous to the CA3 hippocampal field. We directed our tetrodes towards DDi using the data of Giassi et al (2012a), an Atlas of the Gymnotus brain (Corrêa et al 1998) and a high-resolution lab atlas of the *Gymnotus* brain for guidance. This atlas has previously successfully directed tracer injections precisely into DDi (Giassi et al 2012c; Elliott et al 2017). Surface sulci delimiting DD from the more medial dorsomedial pallium and more lateral dorsolateral pallium were used to guide the electrodes to DD and at a site rostral to the very caudal DDmg. The penetration depth was adjusted to pass through DDs but remain confined to DDi (Supplementary Figure 1).

Once the electrode was implanted, the opening of the skull was covered with small pieces of thin plastic sheets (0.5 mil thick FEP film made with Teflon fluoroplastic, CS Hyde Company, Lake Villa, IL). The sheets were then secured to the rest of the skull using a UV curable sealant (Aegis Pit & Fissure Sealant, Keystone Industries, Germany), which was also used to seal any other small remaining openings. After curing with UV light, a mixture of dental cement and Krazy glue was used to further seal the opening and to cover the incision areas on the fish’s skin. At this point the MS-222 solution was slowly diluted with DI water and the PE tubing containing the tetrode was released from the glass capillary by applying a gentle flow of water. Once the fish resumed breathing, it was gently taken out of the setup and transferred to the experimental tank and allowed to recover overnight (Figure 1D).

Recordings were obtained the next day in four out of 5 fish and on day 4 after surgery in one fish that took longer to recover. The recordings were obtained as long as the battery lasted. This was between 5-6 hours in 3 out of 5 fish. For the fish that took a long time to recover (i.e. to resume normal activity level and to swim consistently around the tank) we could only record for about an hour during the 4^th^ day before the battery ended, as we had attempted recording the days before, when the fish was not swimming reliably. For the fifth fish, we have a little over 2 hours of recording after which the fish pulled out the implant and had to be sacrificed right after. At the end of a recording session, we deeply anaesthetized the fish and perfused it in a standard manner (Giassi et al 2012a). We attempted to remove the tetrode and extract the brain with minimal damage and to then section and stain (Cresyl Violet, Giassi et al 2012a) in order to identify the tetrode location. We were successful in locating the tetrode track in two fish (Supplementary Figure 1).

### Experimental setup and trials

The experimental tank was 1.5 m in diameter and fish were tested in shallow water (~10 cm) to facilitate video tracking by restricting fish’s swimming trajectory in 2 dimensions (Figure 1E). A full description of the tank construction and the landmark shapes can be found in (Jun et al 2014a, 2016). The length of the tetrode (15cm) was chosen such that the wireless transmitter float would not exert any force on fish’s head. Landmarks were made of acrylic and were secured to the bottom of the tank using suction cups and could be added or removed. Each trial lasted between 30min – 1hour during which the landmark configuration was stable. Landmarks were sometimes removed or displaced to test the effect on the firing properties of the units. Electric Organ Discharge (EOD) signals were captured using 4 pairs of graphite electrodes (Mars Carbon 2-mm type HB, Staedler, Germany) attached to the tank walls at equal spacing (Jun et al 2014a). All experiments were performed in the dark and the animal’s behavior was monitored under IR illumination using a camera (C910, Logitech, IR filter removed) that was mounted above the tank. The camera acquired images at 1600 × 1200 resolution and had a frame rate of 15 Hz.

#### Wireless data reception and spike sorting

Analog signals received at the receiver were digitized using (CED mkII, Cambridge Electronic Design, UK) and further analyzed using CED’s Spike2 software. EOD signals sensed by four pairs of tank electrodes were also acquired simultaneously using the CED acquisition system. Neural recordings also contained spikes from the EOD. To facilitate spike sorting, EOD spikes were removed from the recording offline, by setting the neural recording trace in the time-window -2.8:2.8 ms around each EOD spike to zero. Figure 2A shows an example recording: the three blue traces are extracellular recordings from the three tetrode channels after removing EOD spikes. and the red trace shows instantaneous EOD rate calculated based on tank electrodes. Spike sorting was done using Spike2 software based on spike waveform shape. The threshold for spike detection was set high and kept constant for all trials in one fish, therefore, only units with high signal to noise ratio were kept for subsequent analysis. Initial sorting by shape was followed by fine tuning spike clusters using principle component analysis and visual inspection of individual spikes (Figure 2B). Due to low firing rate of units, clusters were sometimes not completely separable and therefore some units may be considered as multi-unit.

#### Analysis of EOD rate – spike relationship

The following data analysis were all performed in MATLAB (MathWorks Inc.). Spike-triggered EOD rate (stEODr) and speed (stSpeed) averages were calculated in an 8s window around the timing of each spike (+/- 4 s). Spike-triggered sampling density (stSmpD) was calculated by dividing stEODr by stSpeed for each spike. For each unit then the spike-triggered averages were calculated using all spikes fired by that unit across all trials. The same procedure was repeated 100 times for spikes circularly shifted by a random time (by at least 30 s and at most the length of the trial minus 30 s) for comparison. The average of stEODr over the entire 8 s window for each spike was compared to that calculated for the stEODr for randomly time shifted spikes using Kruskal-Wallis test. To calculate the significance of the peak in stEODr and stSpeed averages, the value of the stEODr and stSpeed for each spike at the time of the peak in the average was calculated. These values were then compared to average EODr and speed calculated over the whole window for all spikes using the Wilcoxon sign rank test. To calculate probability density functions for peak times in stEODr, stSpeed and stSmpD, the timing of the largest peak in EODr, Speed or SmpD was measured for each spike. To calculate the histogram of timings of the dip in stSpeed and stSmpD, the timing of the smallest local minimum was used. The same procedure was repeated for randomly time-shifted spike trains.

#### Analysis of fish’s swimming trajectory and spatial firing rates

Position, and swimming direction of the fish in the experimental tank was calculated using custom software as well as those available from Ty Hedrick’s lab (Hedrick, 2008). To calculated spatial firing properties of the units, the area of the experimental tank was divided into 16 × 16 cm bins and the total number of spikes fired in each bin was divided by the time-spent in that bin. Bins that had less than 5 visits during the whole trial were not included in the analysis. A visit was counted when the fish’s head first arrived at a given bin. If the fish stayed within a bin for more than 1 frame (video frame rate =15 Hz), visit number was still counted as 1, and was allowed to increase only when the fish left the bin and returned to it another time. For visualization purposes color range shown in the firing rate map plots was clipped at 97 percentile of firing rate of that unit across all bins. This was done to avoid bins where the fish spent a very small amount of time to saturate the color plot.

The maximum firing rate per bin is indicated above these plots (Figure 5). Spatial information in bits per spike was calculated using the firing rate maps as described previously (Skaggs et al 1993, Rubin et al 2014):

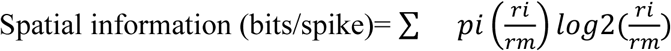

Where p_i_ is the probability of the animal being in the i^th^ bin, calculated as the ratio of time spent in bin i divided by the total trial time, r_i_ is the firing rate in bin i and r_m_ is the mean firing rate for that unit. Only bins with more than 5 visits were included in the analysis. To calculate statistical significance of spatial information, this information measure was recalculated for 1000 randomly time shifted spike trains superimposed on the same swimming trajectory. The spatial information conveyed by a unit was considered significant if it exceeded 95^th^ percentile of the distribution of the information calculated for the randomly shifted spike trains.

To examine the effect of removing a landmark on the firing rate of a given unit we calculated the average firing rate for the bins that were within a 10-cm radius of the landmark, before and after its removal.

#### Swimming direction and landmark location preference analysis

Spike rate during forward (backward) swim was calculated as the number of spikes that occurred during forward (backward) swim divided by the total time spent swimming forward (backward). Because of the small number of spikes, we pooled forward(backward) turns with forward (backward) swims. Direction preference index was calculated as the difference of the firing rate during forward swims and backward swims divided by their sum. Negative values of the direction preference index indicated preference for spiking during backward swims. To calculate the spike-triggered landmark (STLM) matrix, first a 160×120 -element matrix corresponding to the absolute location of landmarks and tank boundary as viewed from the video recording was constructed (each matrix element corresponded to a 10×10 pixels area in the video recording). Matrix elements that contained landmarks or tank boundaries were set to 1 whereas other elements were set to zero. Next, at the time of each spike of a given unit, this matrix was shifted and rotated such that fish’s position was centred in the matrix and its head was facing north. The matrix sum was then calculated for all spikes from a given unit. Similarly, a Position-triggered Landmark (PTLM) matrix was calculated as the sum of such rotated and shifted landmark matrix calculated every 1.33 seconds (the fish’s trajectory, which had a resolution of 15 Hz, was down-sampled 20 times). PTLM matrix elements that were smaller than the lower 10 percentile of all elements were excluded from the analysis. STLM was then normalized to PTLM to calculate the probability of presence of landmarks within 10 cm around the fish. This distance was chosen as a conservative upper range for object detectability with the electric sense based on Jun et al 2016. We then divided the area around the fish into four regions: Left and right side of the fish, each of which was further divided into two regions: anterior 1/3 and posterior 2/3 of the fish’s body. The region located around the anterior third of the fish, corresponds to the regions near fish’s head and upper trunk (gill region) that contains the largest number of electroreceptors (Carr et al 1982; Castello et al 2000). We then calculated a left - right preference index as the difference between the maximum probability on the left side and the right side divided by the sum of the two. We similarly calculated an anterior-posterior preference. Positive left-right (anterior-posterior) preferences corresponded to locations to the right side (anterior) of the fish. For each cell, we then calculated a preference vector with its x and y value equal to the left-right and anterior posterior preference indices, respectively (Figure 7E). For each unit, we used the amplitude and direction of this vector as an indicator of the strength and orientation of preferred landmark location for that unit. Because in all fish the tetrode was implanted on the left hemisphere, landmarks to the left side of the fish were ipsilateral to the location of the electrode and those to the right of the fish were contralateral.

## Acknowledgements

We would like to thank Florian Engert for his helpful comments, suggestions and support for this manuscript. We also thank Bill Ellis for technical support and Erik Harvey-Girard, Armin Bahl, Martin Haesemeyer and Roy Harpaz for their helpful suggestions. This research was supported by NSERC grant 04336 to LM.

---

**Fig S1.**
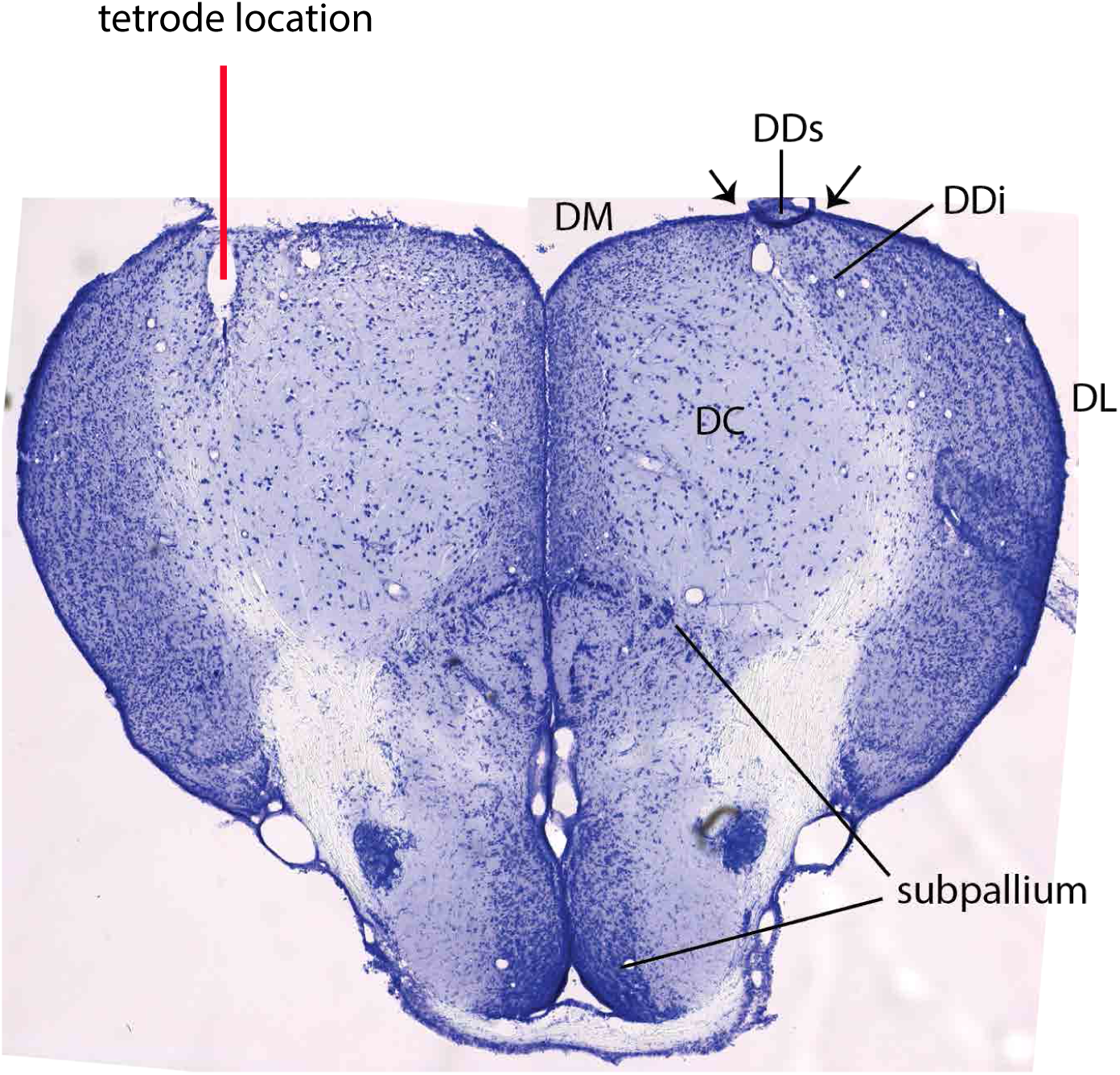
Cresyl violet stained section through the pallium of an implanted fish illustrating the location of the tetrode. Small arrows (right side) indicate the sulci used as an aid in placing the tetrode. Note that the tetrode track passed through the superficial DD (DDs) and ended within DDi. The much lower cell density in DDi compared to DL is evident in this section. DC: central division of dorsal telencephalon DDi: intermediate division of the dorsal portion of dorsal telencephalon (DD, pallium) DDs: superficial division of the dorsal portion of DD DL: dorsolateral pallium DM: dorsomedial pallium

**Fig S2.**
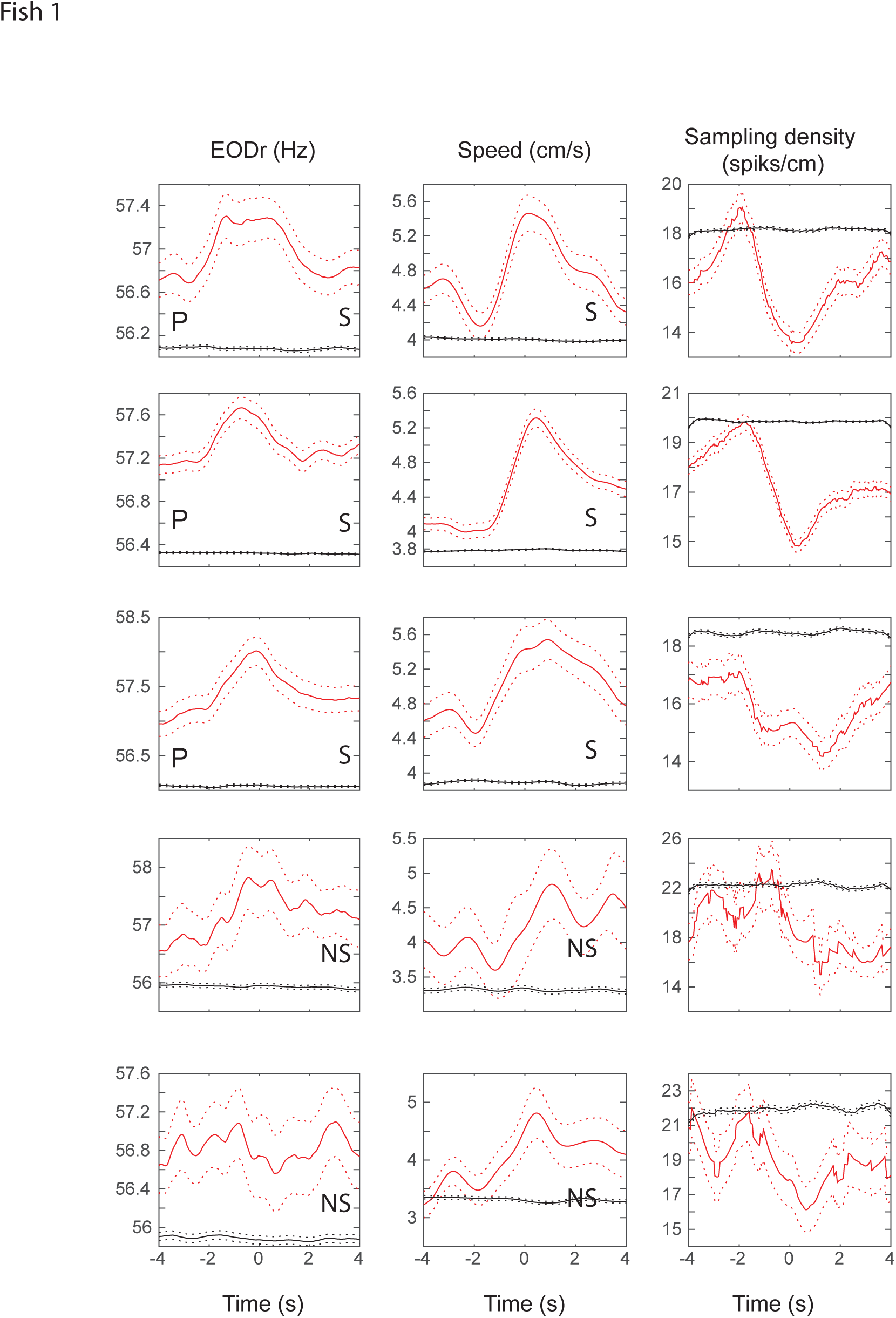
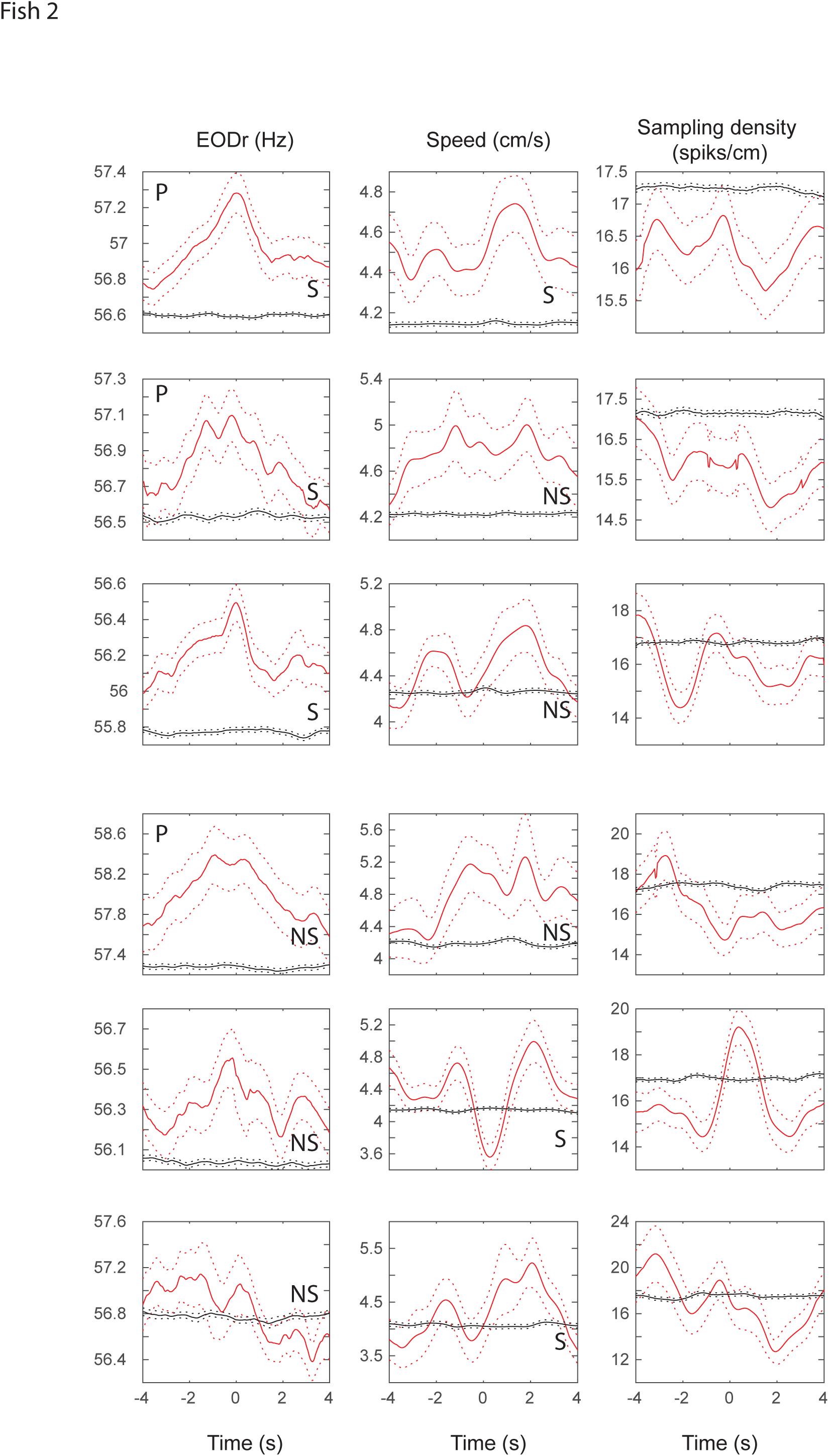
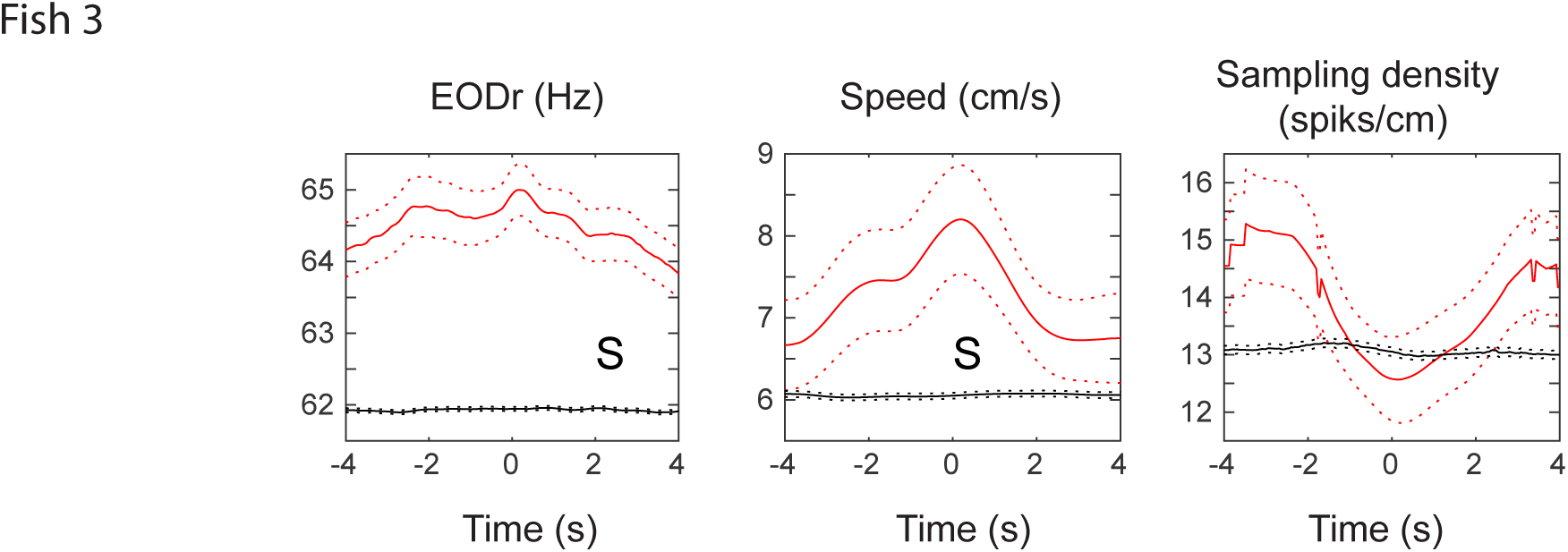
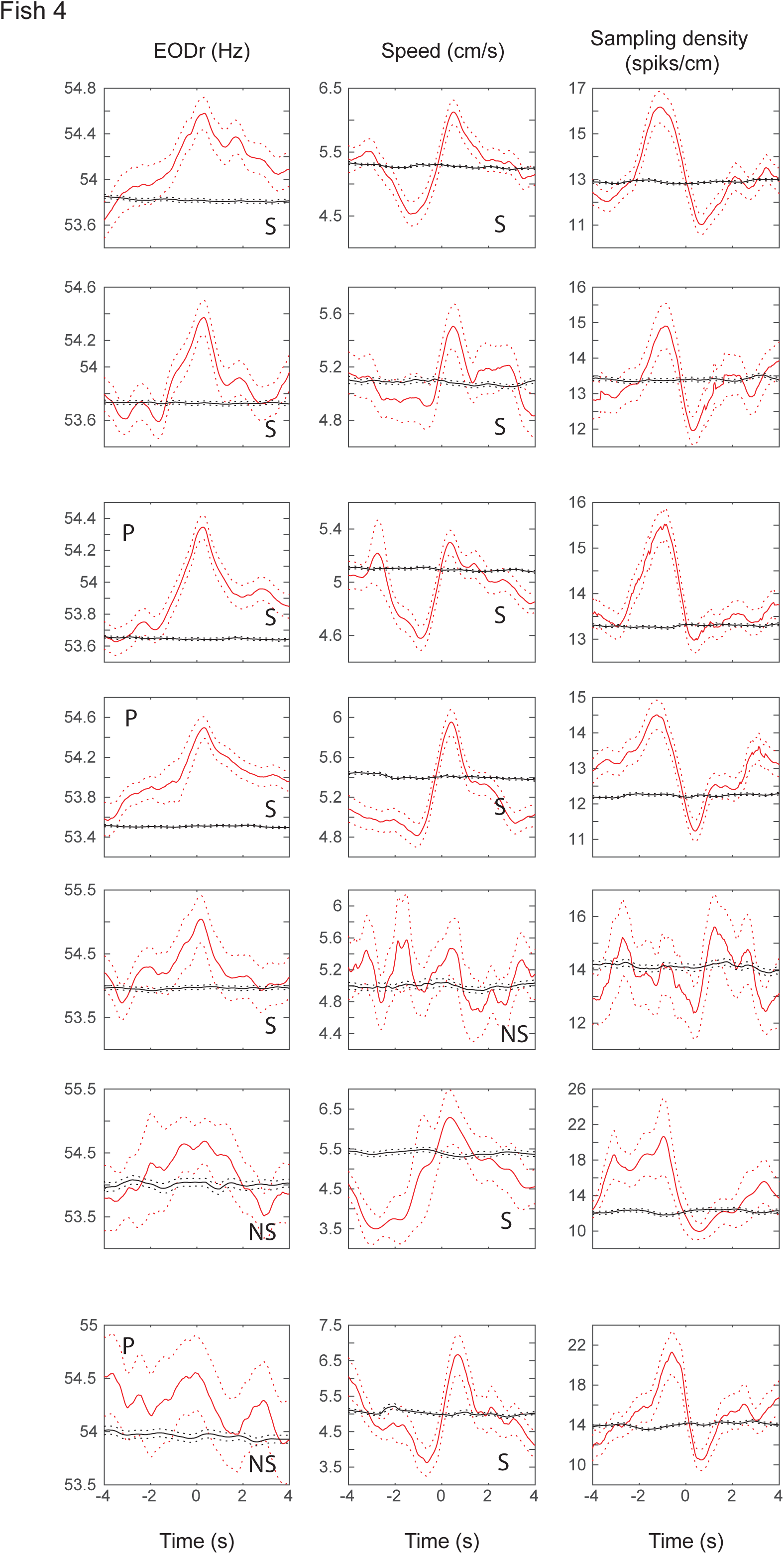
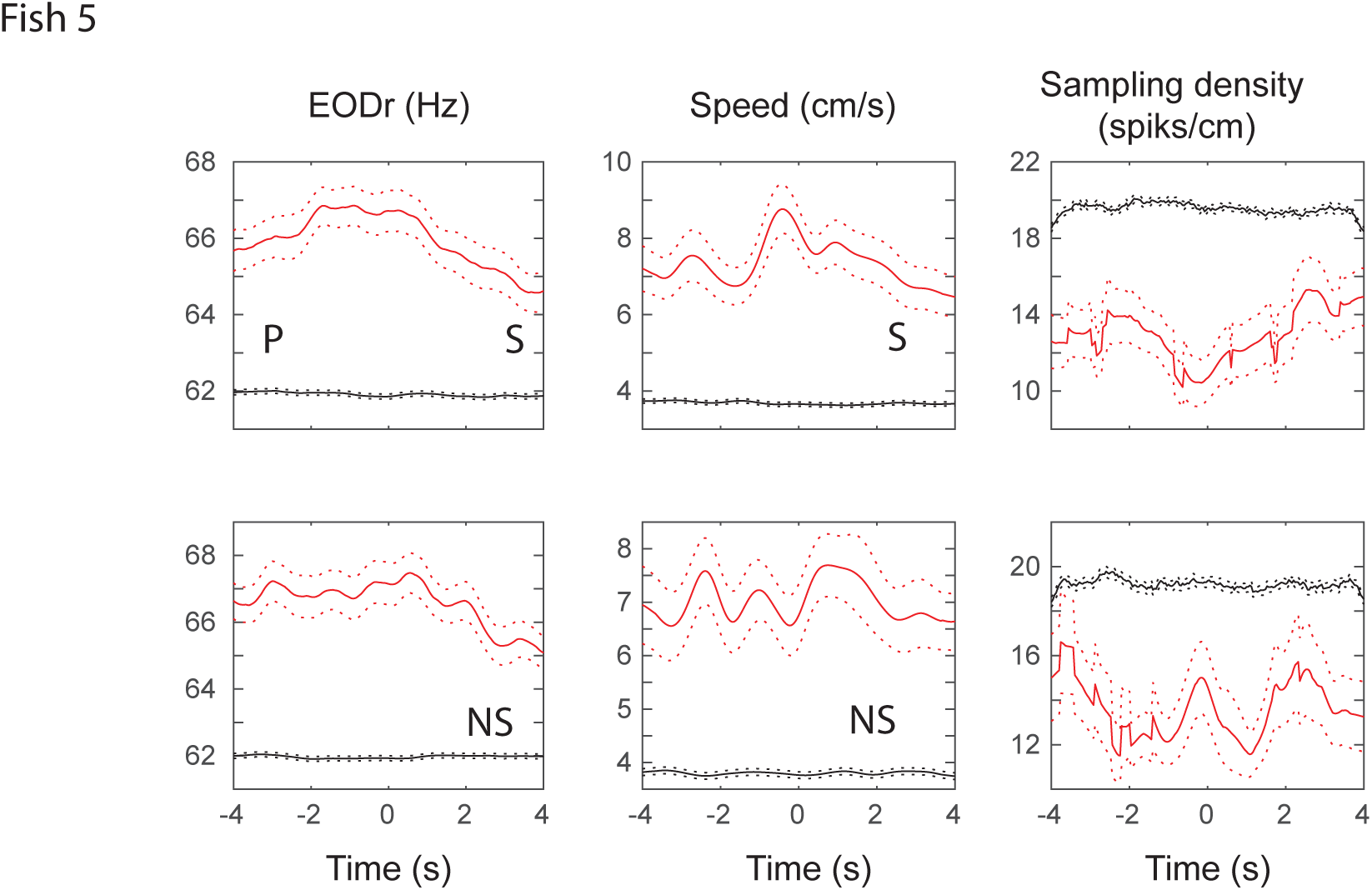
stEOD, stSpeed and stSmpD averages for all units and all fish (red curves). Black curves show the average for 100 random time-shifts of the same spike train. Dashed curves are standard errors. Units with significant (non-significant) peak in their stEOD average are labeled with S (NS). Units that further showed significant place specificity are denoted with a ‘P’.

**Fig S3.**
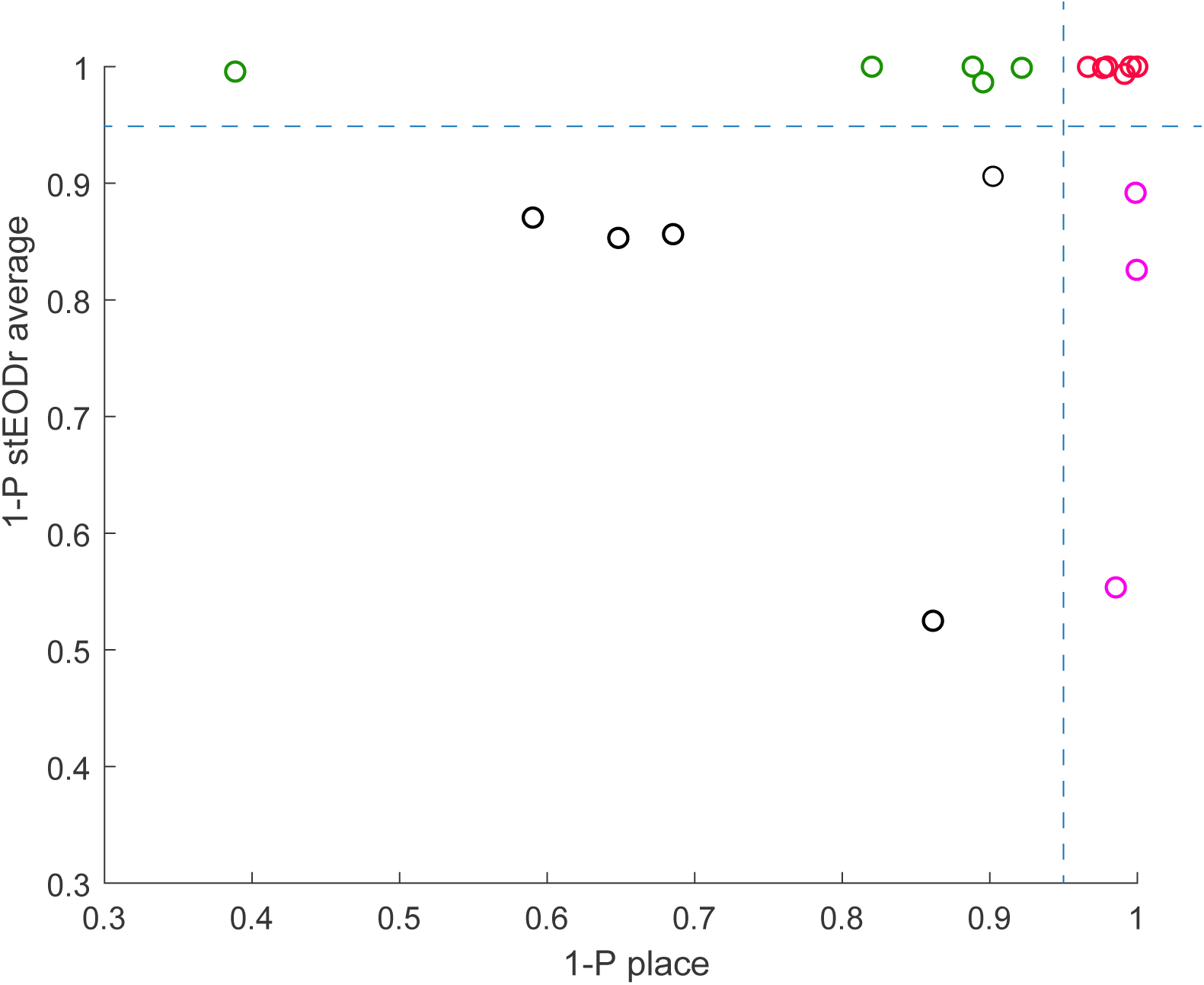
Significance levels for place information and the peak amplitude for the stEODr average for each of the 21 units recorded in 5 fish (1- p values). 11 out of 21 units conveyed significant place information (1-p> 0.95, red and purple circles), 8 of which also had a significant peak in their stEODr average (red circles). Out of the 10 units which did not show place specificity, 5 had significant peaks in their stEODr average (green circles).

## References

Alme CB, Miao C, Jezek K, Treves A, Moser EI, Moser M-B (2014) Place cells in the hippocampus: eleven maps for eleven rooms. PNAS. 111: 18428–18435.

Bastian J (1982) Vision and Electroreception - Integration of sensory information in the optic tectum of the weakly electric fish *Apteronotus-Albifrons*. J Comp Physiol 147: 287–297.

Broglio C, Rodríguez F, Gómez A, Arias JL, Salas C (2010) Selective involvement of the goldfish lateral pallium in spatial memory. Behav Brain Res 210: 191–201.

Eichenbaum H (2017) The role of the hippocampus in navigation is memory. J Neurophysiol 117: 1785–1796.

Elliott SB, Harvey-Girard E, Giassi ACC, Maler L (2017) Hippocampal-like circuitry in the pallium of an electric fish: Possible substrates for recursive pattern separation and completion. J Comp Neurol 525: 8–46.

Elliott SB, Maler L (2015) Stimulus-induced up states in the dorsal pallium of a weakly electric fish. J Neurophysiol 114: 2071–6.

Canfield JG, Mizumori SJY (2004) Methods for chronic neural recording in the telencephalon of freely behaving fish. J Neurosci Methods 133: 127–134.

Carr CE, Maler L, Sas E (1982) Peripheral organization and central projections of the electrosensory nerves in gymnotiform fish. J Comp Neurol 211: 139–153.

Castelló ME, Aguilera PA, Trujillo-Cenóz O, Caputi AA (2000) Electroreception in Gymnotus carapo: pre-receptor processing and the distribution of electroreceptor types. J Exp Biol 203: 3279–3287.

Chen G, King JA, Burgess N, O’Keefe J (2013) How vision and movement combine in the hippocampal place code. PNAS 110: 378–383.

Chersi F, Burgess N (2015) The Cognitive Architecture of Spatial Navigation: Hippocampal and Striatal Contributions. Neuron 88: 64–77.

Corrêa SA, Corrêsa FM, Hoffmann A (1998) Stereotaxic atlas of the telencephalon of the weakly electric fish Gymnotus carapo. J Neurosci Methods 84: 93–100.

Danjo T, Toyoizumi T, Fujisawa S (2018) Spatial representations of self and other in the hippocampus. Science 359: 213–218.

Diamantaki M, Frey M, Berens P, Preston-Ferrer P, Burgalossi A (2016) Sparse activity of identified dentate granule cells during spatial exploration. eLife 2016;5:e20252

Durán E, Ocaña FM, Broglio C, Rodríguez F, Salas C (2010) Lateral but not medial telencephalic pallium ablation impairs the use of goldfish spatial allocentric strategies in a “hole-board” task. Behav Brain Res 214: 480–487.

Fenton AA, Muller RU (1998) Place cell discharge is extremely variable during individual passes of the rat through the firing field. PNAS 95: 3182–3187.

Fenton, AA, Kao, HY, Neymotin, SA, Olypher, A, Vayntrub, Y, Lytton, WW, & Ludvig, N (2008). Unmasking the CA1 ensemble place code by exposures to small and large environments: more place cells and multiple, irregularly arranged, and expanded place fields in the larger space. J Neurosci. 28(44), 11250–62.

Giassi ACC, Harvey-Girard E, Valsamis B, Maler L (2012a) Organization of the gymnotiform fish pallium in relation to learning and memory: I. Cytoarchitectonics and cellular morphology. J Comp Neurol 520: 3314–3337.

Giassi ACC, Duarte TT, Ellis W, Maler L (2012b) Organization of the gymnotiform fish pallium in relation to learning and memory: II. Extrinsic connections. J Comp Neurol 520: 3338–3368.

Giassi ACC, Ellis W, Maler L (2012c) Organization of the gymnotiform fish pallium in relation to learning and memory: III. Intrinsic connections. J Comp Neurol 520: 3369–3394.

Hainmueller T, Bartos M (2018) Parallel emergence of stable and dynamic memory engrams in the hippocampus. Nature 558: 292–296.

Harvey RE, Rutan SA, Willey GR, Siegel JJ, Clark BJ, Yoder RM (2018) Linear Self-Motion Cues Support the Spatial Distribution and Stability of Hippocampal Place Cells. Curr Biol 28:1803–1810.e1805.

Harvey-Girard E, Giassi ACC, Ellis W, Maler L (2012) Organization of the gymnotiform fish pallium in relation to learning and memory: IV. Expression of conserved transcription factors and implications for the evolution of dorsal telencephalon. J Comp Neurol 520: 3395–3413.

Hedrick TL (2008) Software techniques for two- and three-dimensional kinematic measurements of biological and biomimetic systems. Bioinspir Biomim 3: 034001.

Jun, J. J., Longtin, A., & Maler, L. (2014a). Long-term behavioral tracking of freely swimming weakly electric fish. J Vis Exp (85):50962.

Jun JJ, Longtin A, Maler L (2014b) Enhanced sensory sampling precedes self-initiated locomotion in an electric fish. J Exp Biol 217: 3615–3628.

Jun JJ, Longtin A, Maler L (2016) Active sensing associated with spatial learning reveals memory-based attention in an electric fish. J Neurophysiol 115: 2577–2592.

Krahe R, Maler L (2014) Neural maps in the electrosensory system of weakly electric fish. Curr Opin Neurobiol 24: 13–21.

Monaco JD, Rao G, Roth ED, Knierim JJ (2014) Attentive scanning behavior drives one-trial potentiation of hippocampal place fields. Nature 17: 725–731.

Moser EI, Moser M-B, McNaughton BL (2017) Spatial representation in the hippocampal formation: a history. Nature 20: 1448–1464.

Muller RU, Kubie JL (1987) The effects of changes in the environment on the spatial firing of hippocampal complex-spike cells. J Neurosci 7: 1951–1968.

Nelson M, MacIver M (1999) Prey capture in the weakly electric fish Apteronotus albifrons: sensory acquisition strategies and electrosensory consequences. J Exp Biol 202: 1195–1203.

O’Keefe J, Dostrovsky J (1971) The hippocampus as a spatial map. Preliminary evidence from unit activity in the freely-moving rat. Brain Res 34: 171–175.

Ocaña FM, Uceda S, Arias JL, Salas C, Rodríguez F (2017) Dynamics of Goldfish Subregional Hippocampal Pallium Activity throughout Spatial Memory Formation. Brain Behav Evol 90: 154–170.

Omer DB, Maimon SR, Las L, Ulanovsky N (2018) Social place-cells in the bat hippocampus. Science 359: 218–224.

Park E, Dvorak D, Fenton AA (2011) Ensemble Place Codes in Hippocampus: CA1, CA3, and Dentate Gyrus Place Cells Have Multiple Place Fields in Large Environments. PLoS ONE 6:e22349.

Pedraja F, Hofmann V, Lucas KM, Young C, Engelmann J, Lewis JE (2018) Motion parallax in electric sensing. PNAS 115: 573–577.

Poulter S, Hartley T, Lever C (2018) The Neurobiology of Mammalian Navigation. Curr Biol 28:R1023–R1042.

Rodríguez F, López JC, Vargas JP, Gómez Y, Broglio C, Salas C (2002) Conservation of Spatial Memory Function in the Pallial Forebrain of Reptiles and Ray-Finned Fishes. J Neurosci. 22: 2894–2903.

Rolls ET (2016) Pattern separation, completion, and categorisation in the hippocampus and neocortex. Neurobiol Learn Mem 129: 4–28.

Rubin A, Yartsev MM, Ulanovsky N (2014) Encoding of head direction by hippocampal place cells in bats. J Neurosci 34: 1067–1080.

Sarel A, Finkelstein A, Las L, Ulanovsky N (2017) Vectorial representation of spatial goals in the hippocampus of bats. Science 355: 176–180.

Save E, Cressant A, Thinus-Blanc C, Poucet B (1998) Spatial firing of hippocampal place cells in blind rats. J Neurosci. 18: 1818–1826.

Savelli F, Yoganarasimha D, Knierim JJ (2008) Influence of boundary removal on the spatial representations of the medial entorhinal cortex. Hippocampus 18: 1270–1282.

Savelli F, Luck JD, Knierim JJ (2017) Framing of grid cells within and beyond navigation boundaries. eLife 2017;6:e21354

Skaggs WE, McNaughton BL, Wilson MA, Barnes CA (1996) Theta phase precession in hippocampal neuronal populations and the compression of temporal sequences. Hippocampus 6: 149–172.

Solstad T, Boccara CN, Kropff E, Moser MB (2008) Representation of geometric borders in the entorhinal cortex. Science 322: 1865–1868.

Song EY, Kim YB, Kim YH, Jung MW (2005) Role of active movement in place-specific firing of hippocampal neurons. Hippocampus 15(1): 8–17.

Taube JS (2007) The head direction signal: origins and sensory-motor integration. Annu Rev Neurosci 30: 181–207.

Trinh A-T, Harvey-Girard E, Teixeira F, Maler L (2016) Cryptic laminar and columnar organization in the dorsolateral pallium of a weakly electric fish. J Comp Neurol 524: 408–428.

Uceda S, Ocaña FM, Martín-Monzón I, Rodríguez-Expósito B, Durán E, Rodríguez F (2015) Spatial learning-related changes in metabolic brain activity contribute to the delimitation of the hippocampal pallium in goldfish. Behav Brain Res 292: 403–408.

Ulanovsky N, Moss CF (2007) Hippocampal cellular and network activity in freely moving echolocating bats. Nature 10: 224–233.

Wallach A, Harvey-Girard E, Jun JJ, Longtin A, Maler L (2018) A Time-stamp mechanism may provide temporal information necessary for egocentric to allocentric spatial transformations. eLife 2018;7:e36769

Wilson MA, McNaughton BL (1993) Dynamics of the hippocampal ensemble code for space. Science 261: 1055–1058.

Winter SS, Mehlman ML, Clark BJ, Taube JS (2015) Passive Transport Disrupts Grid Signals in the Parahippocampal Cortex. Curr Biol 25: 2493–2502.

